# Dental Caries and Oral Health Behavior Assessment Among Portuguese Adolescents

**DOI:** 10.1101/688911

**Authors:** Nélio Jorge Veiga, Maria Helena de Cecchi, Johnny Martins, Inara Pereira da Cunha, Marcelo de Castro Meneghim

## Abstract

**Introduction:** It is during the school phase that children and adolescents consolidate healthy behaviors, which will contribute to the decrease of diseases, especially in the reduction of dental caries. The main objective of the present study was to assess the decayed, missing and filled deciduous and permanent teeth index and oral health behaviors among Portuguese adolescents.

**Materials and methods:** An observational cross-sectional study was designed including a sample of 694 adolescents between the ages of 12 and 18 years old from five public schools in the Viseu and Guarda districts, Portugal. After a self-administered questionnaire was filled out by the participants, a clinical examination was carried out in order to assess oral status and dental caries identification. A descriptive analysis of the variables was performed using the Chi-square, Mann-Whitney and Kruskal-Wallis tests (p<0.05).

**Results:** The decayed, missing and filled permanent teeth index was 2.91±2.9 and the decayed, missing and filled deciduous teeth index was 1.10±1.4. Of the total sample, 73% consumed sugary food on a daily basis, 54.7% drank bottled water, 50.1% considered oral health good, 70.8% did not report pain in the last 12 months, but noticed gingival bleeding (51.5%). Most adolescents (79.4%) brushed their teeth daily and 60% did not use dental floss. Of the total sample, 96.4% had a dental appointment in the last 12 months, being 46.4% due to prevention treatments. The high decayed, missing and filled deciduous teeth index was associated with low maternal scholarship, male gender and living in a rural residence area (p<0.05). Adolescents who brush their teeth daily presented a good perception about their oral health (p<0.001).

**Conclusions:** Portuguese adolescents presented a low decayed, missing and filled deciduous and permanent teeth index index. The decayed, missing and filled deciduous teeth index was associated with sociodemographic factors. Oral hygiene habits were associated with self-perception of oral health. It is suggested that oral health promotion and prevention programs should be improved in schools in order to reduce the risks of oral disease development.

## Introduction

Dental caries is an ancient disease in the history of mankind being considered today as a public health issue among modern civilization.(1) According to the World Health Organization (WHO) values, for the index of permanent ddecayed, missing and filled permanent teeth (DMFT) at 12 years, Portugal was among the countries with low prevalence of caries with the value of 1.48, reaching the recommended value for the European Region in 2020 (1.50) already in 2006.(2) However, among adolescents with 15 years of age, this index was 3.04.(3) These values, especially among children, are due to the increase in medical-dental treatments resulting from contracting processes with the private sector for the provision of medical-dental care, integrated in the National Program for Oral Health Promotion of the Ministry of Health of Portugal (3). However, there is scarce information in the literature about the distribution of DMFT and its associated risk factors among adolescents, which makes it difficult to develop preventive programs and the organization of health care adapted to the real needs of the population.

Another program of importance for oral health is the National School Health Program, which aims to promote and protect health and prevent disease in the educational community. The program considers that the school has become increasingly prominent in the health of children, adolescents and the rest of the educational community, since it is in school that children and adolescents spend most of their time assimilating new learning and knowledge, transmitting them to the family context.(4)

In this sense, several oral health prevention activities can be implemented in schools to reduce dental caries and other oral diseases, such as discussing the reduction of sugar consumption in the diet, correct tooth brushing, increase young people’s perception of gum bleeding and pain, and educate schoolchildren about the importance of regular dental check-up appointments and the need for fluoride application and use.(5–9)

Both childhood and adolescence are periods of life that represent a greater risk for the development of dental caries, in which health behaviors are consolidated, with emphasis on oral hygiene and eating habits.(10) The lack of healthy lifestyle habits during childhood and adolescence can be an important risk factor for adulthood, and may contribute to serious dental, functional, physical and psychological impairment and consequently reduce quality of life.(11)

The present study consists in the assessment of the decayed, missing and filled teeth index in the deciduous and permanent dentition (dmft and DMFT, respectively) among Portuguese adolescents and identification of the risk factors associated with the cited indexes as well as those associated with oral hygiene habits.

## Materials and methods

The present study is an observational cross-sectional study and obtained approval by the Health Sciences Institute of the Universidade Católica Portuguesa and the formal authorization of the participating schools. The informed and explicit consent of the adolescents participating in the study and their legal guardians was also received. A total of 649 adolescents between the ages of 12 and 18 years old from five public schools in the Viseu and Guarda districts (Aguiar da Beira, Mundão, Abraveses, Mangualde and Satão) during the year 2017 participated in this study. All schools participated in the community oral health program “My Best Smile” developed by Institute of Health Sciences of the Universidade Católica Portuguesa.

Data were collected on socio-demographic situation, eating habits, self-perception of oral health, oral hygiene habits and access to medical-dental services of study participants. For this, the self-applied questionnaire was used, with the following variables:

- Sociodemographic characterization: age, gender, parents’ scholarship and residence area.
- Eating habits: consumption of sugary foods and type of water consumed.
- Self-perception of oral health: how the participants describe oral health, self-reported tooth and gingiva pain, and gingival bleeding during the last 12 months.
- Characterization of oral hygiene habits: daily brushing, mouthwash with fluoride, use of dental floss.
- Access to medical-dental services: dental appointments, frequency, type of service and reason for the dental appointment.

After completing the questionnaire by the participants, the researchers performed the clinical examination using the dmft and DMFT indexes following the criteria of the World Health Organization. The examination was performed by a previously calibrated examiner and scorer. It occurred in a reserved environment at the school, with the adolescent sitting in front of the spotlight and the examiner standing.

Statistical analysis was performed using SPSS-IBM software version 24.0. In the descriptive statistical analysis, absolute and descriptive frequencies were used for variables with nominal measurement level, mean as a measure of central tendency and standard deviation as a measure of dispersion for interval variables (12).

The Chi-square test was used to verify possible associations between dmft and DMFT among the independent variables: age, gender, parents’ educational qualifications and residence area. The association between oral hygiene habits and participants’ description of oral health (self-perception) was assessed.

In the interval variables such as age, it was verified that the criteria of proximity to the normal distribution were not met. For these variables and for ordinal variables, non-parametric statistical tests were used. The Mann-Whitney U was used to test differences between two groups, for three or more groups the Kruskal-Wallis test was used.

## Results

The sample presented an average of 13.9 years of age. Of the 694 participants, 360 (51.9%) belonged to the male gender and 334 (48.1%) to the female gender (table 1).

**Table 1.** Descriptive statistics of the sample studied, by gender.

| Gender | n | Age |  |  |  |  |
| --- | --- | --- | --- | --- | --- | --- |
|  |  | % | Mínimum | Maximum | Average | Standard deviation |
| Male | 360 | 51.9 | 12 | 18 | 13.8 | 1.8 |
| Female | 334 | 48.1 | 12 | 18 | 14.0 | 1.8 |
| Total | 694 | 100.0 | 12 | 18 | 13.9 | 1.8 |

On average, the sample had 1.95 decayed teeth. Two hundred and eighty-one adolescents had 1-2 decayed teeth (29.2%) and 252 adolescents had no decayed teeth (39.3%). Lost teeth presented an average of 0.17. Most of the adolescents had not lost any teeth (n=574, 89.5%). The filled teeth had a mean of 0.81. Most of the adolescents had no filled teeth (n=418, 65.2%). The DMFT showed an average of 2.91. The DMFT score was one to three for 36.9% of the sample (table 2).

**Table 2.** DMFT and dmft scores among the studied sample.

| DMFT and dmft | Minimum | Maximum | Average | Standard deviation |
| --- | --- | --- | --- | --- |
| Decayed permanent teeth (n=641) | 0 | 12 | 1.95 | 2.3 |
| Missing permanent teeth (n=641) | 0 | 7 | 0.17 | 0.6 |
| Filled permanent teeth (n=641) | 0 | 11 | 0.81 | 1.4 |
| DMFT (n=645) | 0 | 15 | 2.91 | 2.9 |
| Decayed deciduous teeth (n=157) | 0 | 7 | 1.22 | 1.7 |
| Missing permanent teeth (n=157) | 0 | 4 | 0.17 | 0.6 |
| Filled permanent teeth (n=157) | 0 | 5 | 0.17 | 0.7 |
| dmft (n=157) | 0 | 7 | 1.10 | 1.4 |

Regarding deciduous dentition, caries was on average 1.22 (n=157), with 49% of the adolescents with no decayed tooth. In the lost primary teeth, a mean of 0.17 was observed. Most of the adolescents observed had no deciduous tooth lost (n=144; 91.7%). The majority of the adolescents had no filled teeth (n=143; 91.1%). The DMFT presented an average of 1.10. The DMFT score ranged from zero to 51% (n = 80) of the sample (table 2).

Table 3 shows that most adolescents reported consuming sugary foods at times (n=316, 73%), and reported consuming bottled water (n=169, 54.7%). In oral hygiene habits, half of the adolescentes considered having “good” oral health (n=218, 50.1%). Most of the adolescents did not present dental pain in the last 12 months (n=305, 70.8%). Regarding gingival hemorrhage or gingival pain, the adolescents did not present these symptoms (n=210; 48.5%).

**Table 3.**
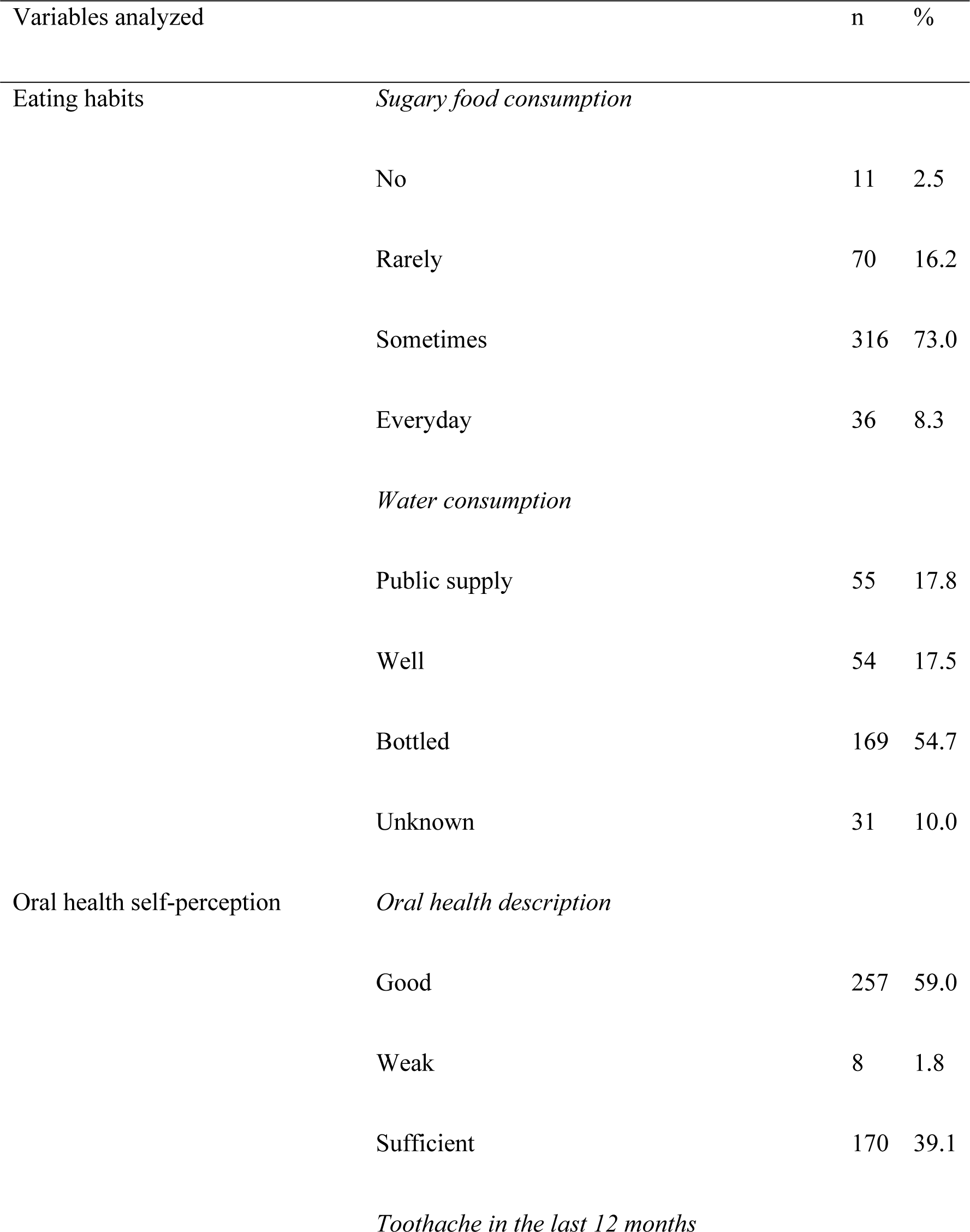
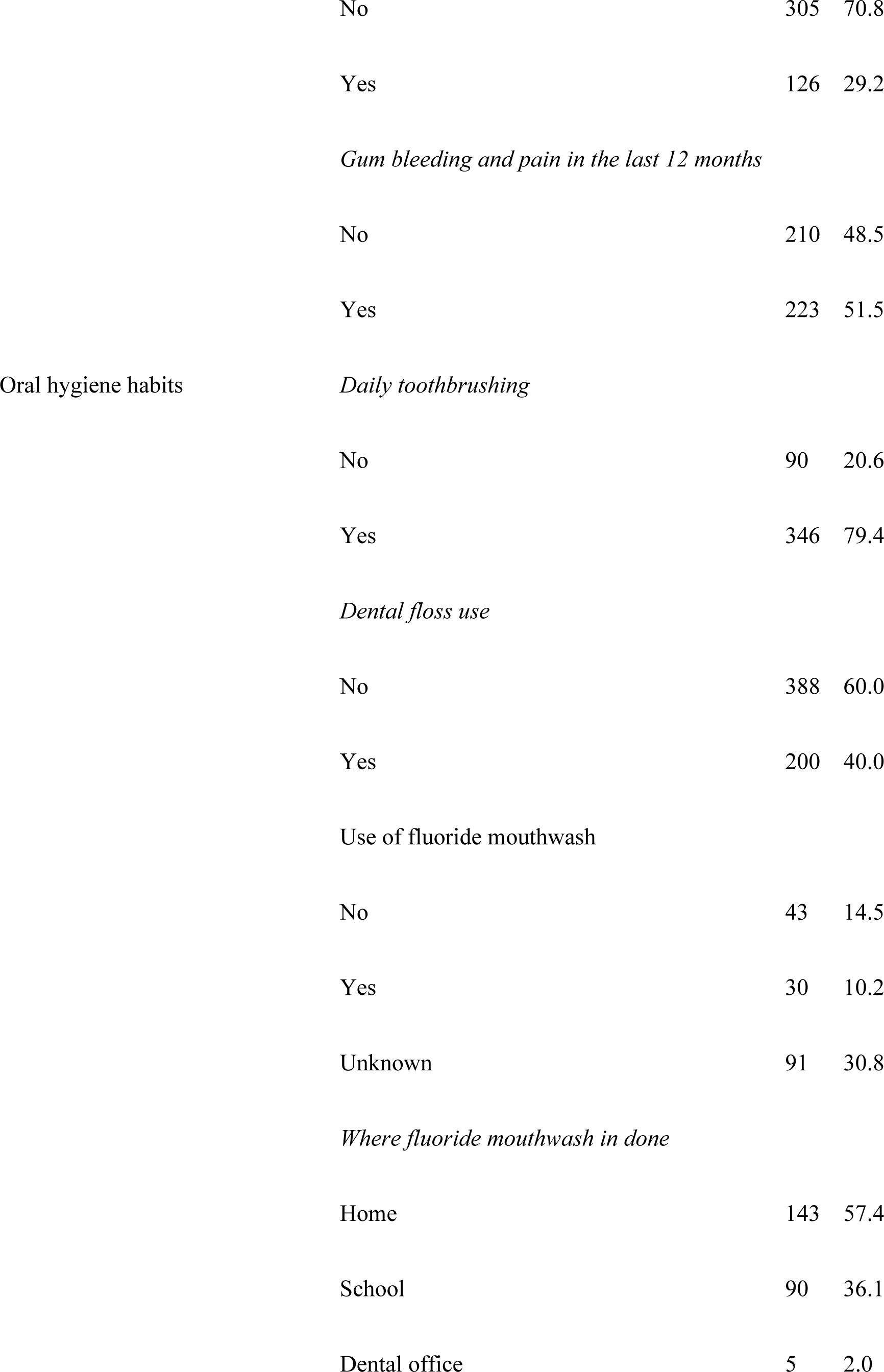
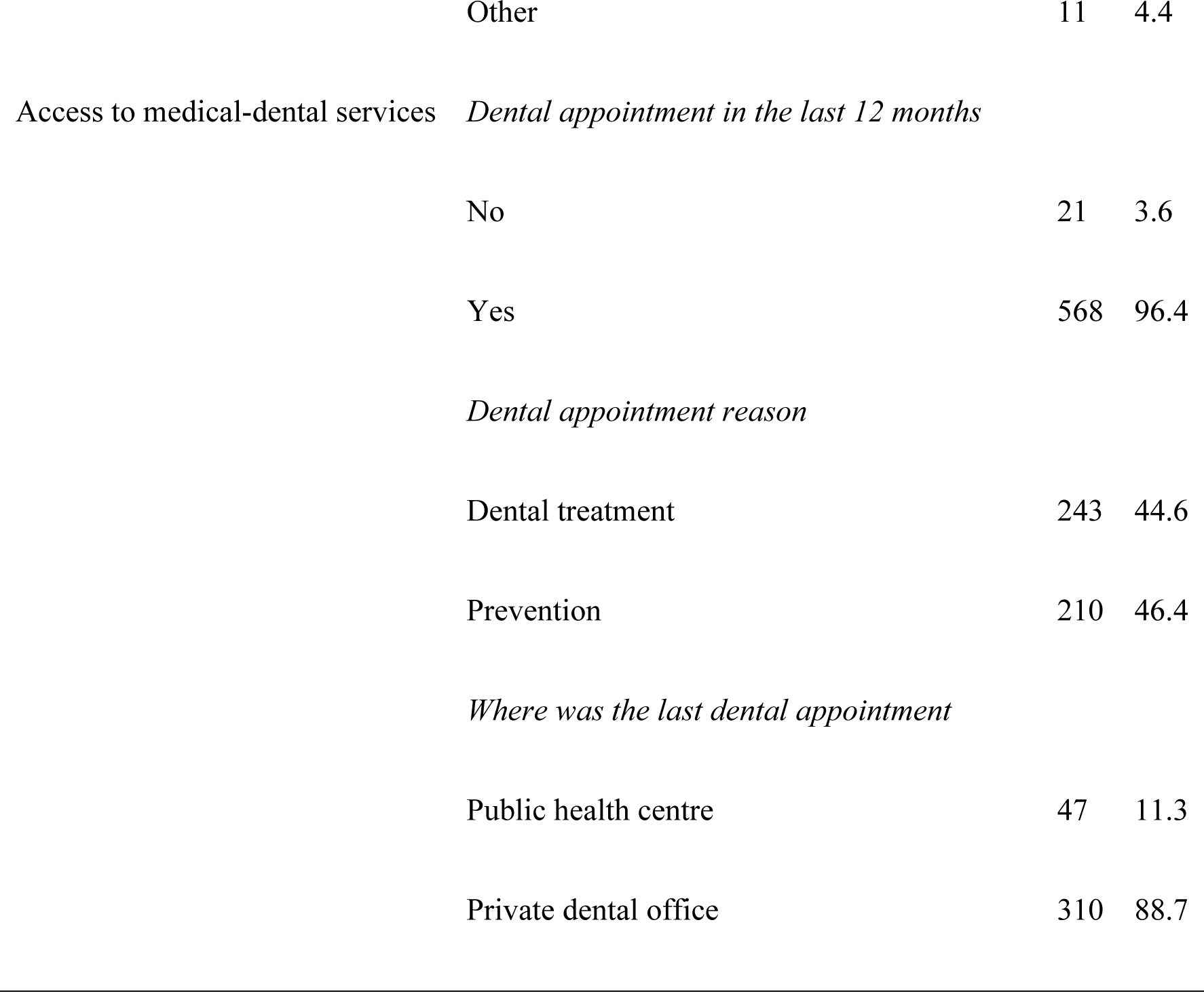
Distribution of the frequency of eating habits, self-perception of oral health, oral hygiene habits and access to dental services among the adolescents.

Regarding oral hygiene habits, most adolescents (n=346, 79.4%) stated that they brush their teeth every day, while 90 (20.6%) adolescents do not brush their teeth daily, and 388 (60%) do not use dental floss. Regarding fluoride mouthwash, the majority of the adolescents affirmed to perform this action (72.1%). When questioned about where they performed mouthwash, the majority of the adolescents answered that they did at home (n=143; 57.4%) (table 3).

Still on table 3, with regard to access to medical-dental services, 361 (61.4%) had a dental appointment during the last 12 months. The most frequent reason for a dental appointment was curative dental treatment (n=244, 44.6%). The dental appointment site occurred mainly in the private dental office (n=370; 88.7%).

In table 4, the DMFT did not obtain statistically significant associations with sociodemographic factors. For dmft, there was a statistically significant correlation between the scholarship of the mother, male gender and residence area (p <0.05).

**Table 2.**
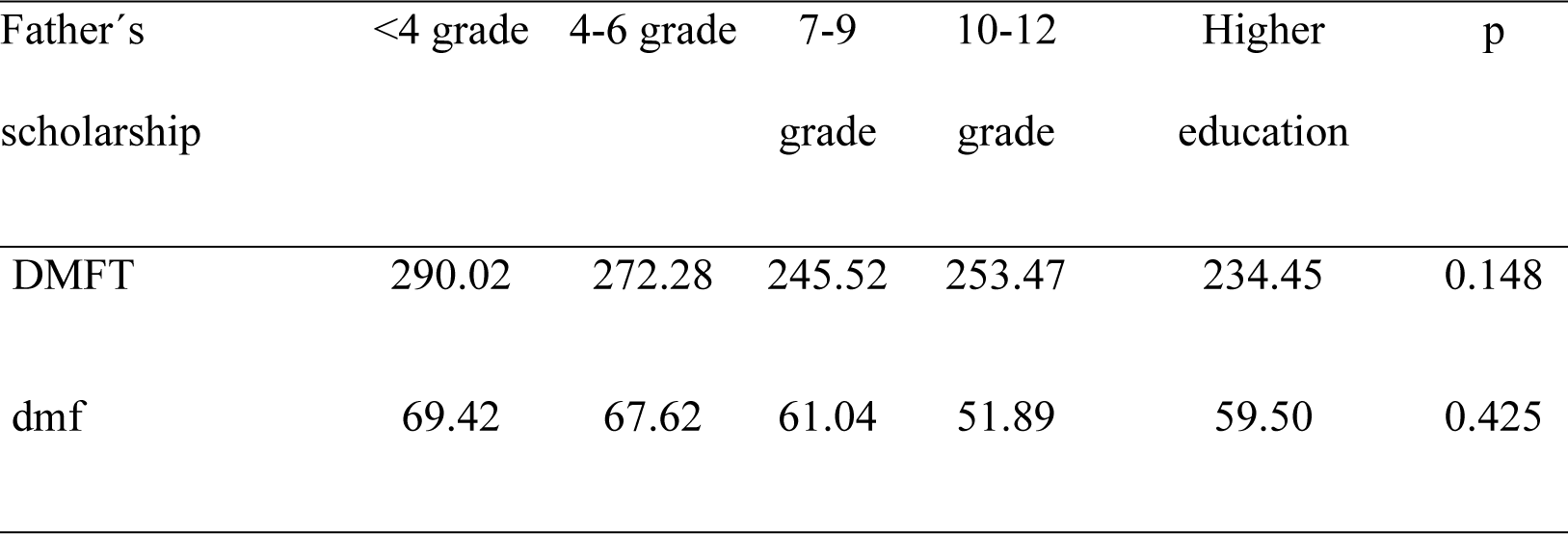
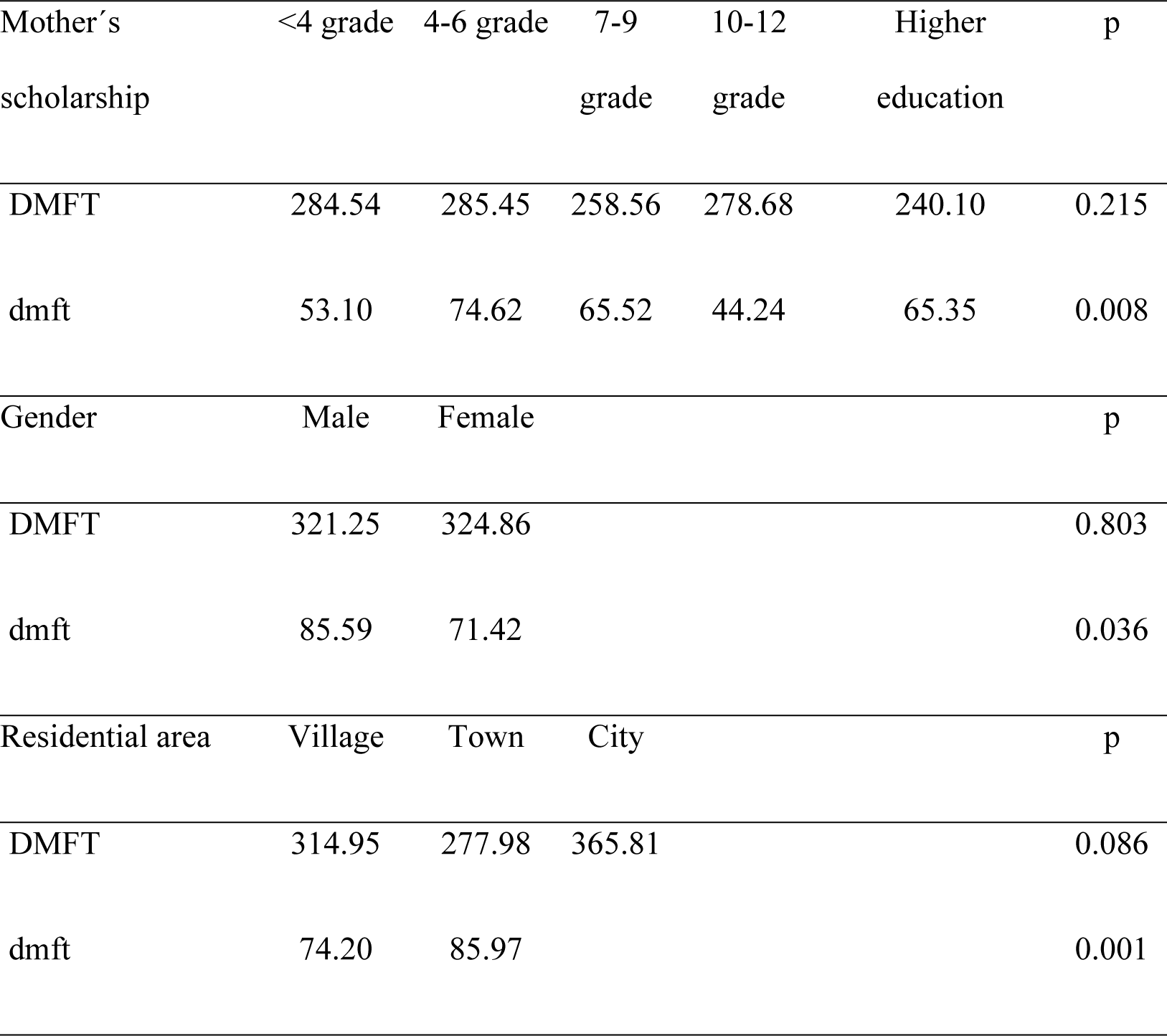
Association between parents’ scholarship and adolescents’ DMFT and dmft (presented in average scores).

Adolescents who brush their teeth every day have a more positive perception of their oral health than subjects who do not (p<0.001). No differences were observed between those using and not using dental floss in the perception about their oral health (p=0.565). The perception of oral health is not correlated with the number of daily brushings (p=0.132).

## Discussion

In the present study there were no differences in distribution among male and female adolescents. Regarding the DMFT index, this was higher than that found in the National Study of Prevalence of Oral Diseases published in 2008, in Portugal (DMFT index=1.48). However, when compared to another study, such as that of Barata *et al.*, the DMFT index was lower.(13) This is because, Barata *et al.*, found an average DMFT index of around 4.05 among Portuguese students, and the presence of caries in 71.8% of the participants.(13) In Brazil, a study also performed with adolescents showed an average DMFT index of 3.28.(5)

Adolescents presented 1.95 decayed teeth, ranging from 0 to 12 teeth. In addition, 35.7% of the sample presented a greater burden of disease (4 or more decayed teeth). These data are important because it identifies a group of adolescents with a higher number of dental caries, and another one in which the disease is very little or totally absent, evidencing a possible polarization. According to Peres *et al.*, the decline of dental caries in several countries was accompanied by the phenomenon of polarization, which consists in the concentration of most diseases and treatment needs in a small part of the population. (14) We have moved from a situation of high disease prevalence to a scenario in which a large proportion of individuals are found to be free of dental caries. The non-uniform distribution also occurs in several studies and is associated with social inequalities (15), since the situation is more severe in the economically disadvantaged and low-grade classes. Regarding the dmft index, there was an average of 1.10 deciduous decayed teeth ranging from 0 to 7 deciduous decayed teeth, among 69% of the sample. Despite the homogeneous situation of caries distribution in this segment, it is necessary to maintain attention for preventive care. Comparing these results with previous studies, also performed in the region of Viseu, it is observed that there was a higher prevalence of caries in the results found. Amaral *et al*., in a study with 76 children, identified the presence of decayed teeth in only 15.3% of the sample.(16)

The experience of caries among 12-year-old adolescents is strongly associated with those with less privileged socioeconomic conditions, whose parents, especially mothers, have a lower educational level and there are strong indications of the association between the disease and the male gender.(17–19) The residence area is another factor that contributes to the prevalence of dental caries.(20) This information corroborates with the present study, which found a significant association between the mother’s scholarship, adolescents’ gender and residence area for the dmft variable. In addition to social factors, it is known that the dental caries index is related to the subjects’ behavioral aspects. Thus, in order to think about preventive actions, it is necessary to understand what care is taken in the daily activities of the adolescents (21).

In this sense, some of the measures to prevent the prevalence of dental caries were investigated in the present study, such as sugar consumption. The frequency of consumption of foods with sucrose has been constantly associated with the prevalence of dental caries among younger people.(5,22) However, not always adolescents who consume higher amounts of cariogenic foods develop caries, since the manifestation of caries is dependent on the time of exposure and other primary and secondary etiological factors. The present results showed that 73% of adolescents stated that they consume sugar daily, a pattern considered negative for food, and that it may contribute to other health problems such as obesity and diabetes.(16) Regarding the association between eating habits, it is also important to investigate the consumption of fluoridated water, since this is an effective preventive measure against dental caries (23). About this, most adolescents reported consuming bottled water, but they do not recall the use of fluoride in the daily mouthwash.

Oral hygiene habits are essential in the removal of plaque and food debris.(21,24) According to the Health General Directory of Portugal, in 2008, approximately 50% of children aged 6 years and around 69% of young people aged 12 and 15 have the daily habit of toothbrushing.(3) To confirm this information, a study carried out in Portugal found that 23.5% of a sample of 7644 adolescents brush their teeth twice a day, and only 4.4% use dental floss.(25) The present study identified a greater frequency in the use of the toothbrush compared to the use of the dental floss. The use of dental floss prevents the development of dental caries on the interproximal surfaces of the teeth and periodontal diseases, and its use is recommended, making it essential to plan educational actions for its use.(26,27)

For dental appointments, they should be regular or at least once every six months, which contributes to the early and immediate diagnosis of oral diseases, information on the most appropriate treatments and application of preventive measures, such as topical application of fissure sealants and fluorides.(28) It has been identified in studies that Portuguese young people frequently have a dental appointment (in the last 12 months), and most are check-up dental appointments.(25) This result is interesting because they may be the result of public health policies that are promoting greater access to the Portuguese population for medical-dental health care. There is a tendency in the literature to assess the subject’s own perception of their oral health. Self-perception of the oral condition has been used as an indicator of individuals’ behavior regarding the search for medical-dental treatments.(29) There is evidence that people who identify a good oral health condition have a lower prevalence of dental caries.(29) Thus, it is plausible the association found between good oral health habits and the good perception of oral health among adolescents.

## Conclusions

The results obtained in the present study were consistent with those described in the literature and indicate that actions to control dental caries among adolescents should be strengthened, and especially aimed at the development of more favorable and efficient oral health behaviors. The results also demonstrated that a better oral hygiene habit, such as frequent toothbrushing, is associated with a more positive perception of oral health. Portuguese adolescents presented a low DMFT and dmft index and are was associated with sociodemographic factors. Oral hygiene habits were associated with self-perception of oral health. It is suggested that oral health promotion and prevention programs should be improved in schools in order to reduce the risks of oral disease development.

## Acknowledgments

The authors are deeply indebted to the teachers and adolescents of the School Groups for the participation and important contribution to this study.

## Ethical declarations

The study was approved by the Health Sciences Institute of the Portuguese Catholic University and obtained formal authorization from the participating schools. The study was in accordance with the 1964 Helsinki Declaration on Ethics Approval and consent to participate in research. The Free and Informed Consent Terms were obtained from the parents before starting the study.

## Financing source

The authors declare that the study had no sources of funding.

## Author Contributions

Conceived and designed the experiments: NJV JM. Performed the experiments: NJV JM. Analyzed the data: NJV JM MHC. Contributed reagents/materials/analysis tools: NJV MHC JM IPC. Wrote the paper: NJV MHC JM IPC MCM.

## References

1. Carounanidy U, Sathyanarayanan R. Dental Caries - A complete changeover - (Part I). J. Conserv Dent. 2009;12(2):46–54.

2. World Health Organization. Oral heath surveys: basic methods. 4th ed. Geneva: World Health Organization; 1997

3. General Directory of Health of Portugal. National Study of Prevalence of Oral Diseases. Program for the Promotion of Oral Health in Schools. 2008; Lisbon.

4. General Directory of Health of Portugal. Program for the Promotion of Oral Health. [cited 2019 Feb 4]. Available from: https://www.dgs.pt/paginas-de-sistema/saude-de-a-a-z/programa-nacional-de-promocao-de-saude-oral.aspx.

5. Rodrigues MA, Silva RP, Pereira PF. Relationship of dental caries with nutritional status, social factors and behavioral in adolescents from 15 to 19 years. RASBRAN – Rev Associ Bras Nutri. 2018;9(2):103–10.

6. Silva Junior IF, Aguiar NL, Barros RC, Arantes DC, Nascimento LS. Teenager’s Oral Health: Literature Review. Rev Adolesc. Saúde. 2016;13 13(Supl. 1):95–103.

7. Araújo MVA, Barriga ALC, Emmi DT, Pinheiro HHC, Barroso RFF. Caries prevalence, autoperception and impact on oral health in teenagers in Marajó island - PA. RDAPO: Rev Digi Acad Paraense de Odontologia Belém-PA. 2017;1:11–17.

8. Martins MMF, Aquino RA, Pomponet ML, Pinto-Júnior EPP, Amorim LDAF. Adolescent and youth access to primary health care services in a city in the state of Bahia, Brazil. Cad. Saúde Pública. 2019;35(1):e00044718.

9. Antunes J, Toporcov T, Bastos J, Frazão P, Narvai P, Peres M. Oral health in the agenda of priorities in public health. Rev Saúde Pública. 2016;50:57.

10. Almeida TF, Cangussu MCT, Chaves SCL, Amorim TM. Oral health status of children, adolescents, and adults registered in Family Health Units Service in the Municipality of Salvador, State of Bahia, Brazil, in 2005. Epidemiol Serviço de Saúde. 2013;21(1):109–18.

11. Barros CMSB. Technical manual on oral health education. Rio de Janeiro: SESC, National Department. 2007;53–5.

12. Pestana M, Gageiro J. Data analysis for social sciences: the complementarity of SPSS. 4. ed. Lisbon: Edições Silabo, 2005.

13. Barata C, Veiga N, Mendesc C, Araújo F, Ribeiro O, Coelho I. DMFT and oral health behaviours assessment in a sample of adolescents of Mangualde. Rev Port Estomatol Med Dent Cir Maxilofac. 2013;54(1):27–32.

14. Peres SHCS, Carvalho FS, Paz CC, Magalhães JB, Pereira LJR. Polarization of dental caries in teen-agers in the southwest of the State of São Paulo, Brazil. Ciênc. saúde coletiva. 2008;13(Suppl 2):2155–62.

15. Da Costa AM, Tôrres LHN, Meirelles MPR, Cypriano S, Batista MJ, Sousa MLR. Low prevalence of caries: polarization group and the importance of family aspects. Rev Odontol Bras Central 2016;25(72):6–11.

16. Amaral A, Melão N. Health profile of children monitored in primary care consultation in Viseu, Portugal. Rev Port Saúde Pública. 2016;34(1):53–60.

17. Souza ME, Pereira SM, Castilho ARF, Pereira LJ, Pardi V, Pereira AC. Relationship among socioeconomic and clinical factors with oral health, in schoolchildren from rural areas: a longitudinal study. RFO UPF 2015;20(2):208–15.

18. Costa MM, Souto IC, Barroso KM, Paredes SO. Factors associated with the experience of dental caries in public schools at a small city in the Northeast of Brazil Rev. Bras. Pesq. Saúde, Vitória, 2017;19(3):32–40.

19. Reis HC, Pontres IC, Furlanetto DLC, Amaral LD, Castro-filho AA. Epidemiological Survey of Dental Caries in Schoolchildren of 2 Schools of the Public Network of the Federal District. Oral Sci. 2013;5(1):5–8.

20. Engelmann JL, Tomazoni F, Oliveira MDM, Ardenghi TM. Association between Dental Caries and Socioeconomic Factors in Schoolchildren - A Multilevel Analysis. Braz. Dent. J. 2016;27(1):72–8.

21. Daniel S, Harfst S, Wilder R. Mosby’s dental hygiene: concepts, cases and competencies. 2^a^ Ed. St Louis: Mosby Elsevier. 2008;440–55.

22. Bonotto DMV, Pintarelli TP, Santin G, Montes GR, Ferreira FM, Fraiz FC. Dental caries and gender in adolescentes. RFO 2015;20(2):202–7.

23. Gonçalves MM, Leles CR, Freire MCM. Associations between Caries among Children and Household Sugar Procurement, Exposure to Fluoridated Water and Socioeconomic Indicators in the Brazilian Capital Cities. Inter J Dent. 2013;2013:492790.

24. Hattne K, Folke S, Twetman S. Attitudes to oral hearth among adolescents with high caries risk. Acta Odont Scand. 2001;65:206–213.

25. Pereira C, Veiga N, Amaral O, Pereira J. Oral health behaviors among portuguese adolescents. Rev Port Saúde Pública. 2013;31(2):145–52.

26. Pereira A. Odontologia em Saúde Colectiva - Planejando acções e promovendo saúde. 1^a^ Ed. Porto Alegre: Artmed Editora; 2003.

27. Sambunjak D, Nickerson J, Poklepovic T, Johnson T, Imai P, Tugwell P, et al. Flossing for the management of periodontal diseases and dental caries in adults. Cochrane Database Syst Rev. 2011 Dec 7;(12):CD008829.

28. Crocombe L, Broadbent J, Thomson W, Brennan D, Poulton R. Impact of dental visiting trajectory patterns on clinical oral health and oral health-related quality if life. J Public Health Dent. 2012;72:36–44.

29. Gibilini C, Esmeriz CEC, Volpato LF, Meneghim ZMAP, Dias SD, Sousa MLR. Access to dental services and self-perception of oral health in adolescents, adults, and the elderly. Arq. Odontol. 2010;46(4):213–23.

